# Divergent interactions maintain the quaternary octameric structure of a new family of esterases

**DOI:** 10.1101/466904

**Authors:** Onit Alalouf, Rachel Salama, Ofir Tal, Noa Lavid, Shifra Lansky, Gil Shoham, Yuval Shoham

## Abstract

Protein oligomerization contributes significantly to the stability and function of enzymes, and the interacting interfaces that create the oligomers are expected to be conserved. The acetyl-xylo-oligosaccharide esterase, Axe2, from the thermophilic bacterium *Geobacillus stearothermophilus* represents a new family of esterases belonging to the SGNH superfamily of hydrolytic enzymes, and has a unique doughnut-like homo-octameric configuration, composed of four homo-dimers. The dimers of Axe2 are held together mainly by clusters of hydrogen bonds involving Tyr184 and Arg192, as was demonstrated by site directed mutagenesis. Dimeric mutants obtained by single amino acid replacements were inactive towards 2-naphthyl acetate, indicating the necessity of the octameric assembly for catalysis. The crystal structure of two homologous proteins (PDB 3RJT and 5JD3) reveal the same tertiary fold and octameric ring structure as of Axe2. Surprisingly, these octameric structures appear to be maintained by different sets of amino acids involving Asn183 in 3RJT and His185 in 5JD3 instead of Tyr184 in Axe2. These findings prompt us to investigate five more homologues proteins, which were found to have similar octameric structures, despite significant changes in their key residues. We revealed a conserved quaternary structure, which is maintained *via* non-conserved interactions.

## Introduction

Protein oligomerization is a widespread phenomenon in which 70-80% of all proteins are composed of a number of subunits, connected through non-covalent bonds to form functional oligomeric proteins. Many studies demonstrate that oligomerization contributes to protein stability, activity, affinity and specificity (1) (2) (3) (4) (5) (6). Oligomers vary in complexity, and may consist of identical monomers (homo-oligomers), or two or more different types of subunits (hetero-oligomers). Homo-oligomerization is the most common form among proteins, and homo-dimers are the most prevalent (3) (7). The frequency of higher order of homo-oligomers decreases with the increase in number of subunits. Homooligomer symmetry is either isologous or heterologous. Isologous assemblies involve association of identical interfaces and hence the same contact residues of both subunits, which are related by a two-fold symmetry axis. This type of association may result in a duplication of interaction at the same interface. Heterologous assemblies involve association of non-identical interfaces (2). A study based on a large data set of protein associations concluded that most interfaces show a significantly higher level of sequence conservation than the remainder of the protein surface (8). As data accumulates, it appears now that the interface sites have been conserved throughout evolution due to their structural importance (9) (10) (11). Analysis of conservation patterns is a useful tool for prediction of protein interfaces (12) and for evolutionary models (13), however, conservation by itself is not sufficient (14).

The intracellular Axe2 enzyme from *Geobacillus stearothermophilus* is a xylo-oligosaccharide serine esterase. The enzyme hydrolyzes ester linkages of acetyl groups at positions 2 and/or 3 of the xylose moieties, facilitating the utilization of plant-derived polysaccharides by the bacterium (15) (16). Axe2 belongs to the Lipase GDSL family (Pfam accession PF00657) (17) (18), part of the SGNH superfamily of hydrolytic enzymes, and represents a new family of esterases (15). The crystal structures of Axe2 (PDB 4JHL and 3W7V) and its mutants (PDB 5BN1, 4OAO, 4OAP, 4JJ4, 4JJ6, 4JKO) were recently determined (19). The Axe2 monomer corresponds to the SGNH hydrolase fold, consisting of five central parallel β-sheets, flanked by two layers of helices (18) (20). As in the case of other SGNH family members, the catalytic triad is made of His and Asp residues situated on a catalytic loop located between two α helices, and of a Ser residue situated on an adjacent loop (21). The His residue acts as a general base and increases the nucleophilicity of the Ser. The Asp residue is assumed to be involved in the stabilization of the ion-pair generated between the imidazoliun ion and the negatively charged intermediate, and may has a role in maintaining a correct orientation of the His relative to the Ser (22). The quaternary structure of Axe2 is a “doughnut-shaped” homo-octamer made of four staggered dimers (**Fig. 1A**, left). Several cases of hexameric doughnut-shaped hydrolases have been reported in the past, but Axe2 appears to be the first hydrolase with an octameric doughnut-shaped assembly possessing inward-facing active sites. This special octameric assembly probably contributes to the overall stability of the enzyme (19) (6). In the present study, we first determined the key interactions that maintain the octameric structure of the Axe2-WT protein. We then investigated the corresponding interactions in Axe2 homologues from *Alicyclobacillus acidocaldarius* (PDB 3RJT) and from the metagenome of Lake Arreo, Spain (PDB 5JD3), which have similar (almost identical) quaternary structures to Axe2 (**Fig. 1A**, center and right, **Table 1**). To our surprise, we found that the three proteins maintain their octameric structures *via* different interactions. These findings prompted us to investigate the quaternary structures of five additional Axe2 homologues. Since the crystal structures of these homologues are not available, we used alternative techniques to determine their functional protein assembly, their shape, and the potential residues that take part in their protein-protein interactions. We demonstrate that all of these Axe2 homologues appear as protein octamers, most likely in a quaternary assembly similar to Axe2, 3RJT, and 5JD3. Our results indicate that these conserved quaternary structures are maintained by inter-subunit interactions involving a non-conserved residues. Hence, we demonstrate, for the first time to our knowledge, a case of sequence divergence in maintaining similar protein quaternary structures.

**Table 1:** Calculated r.m.s.d (PyMol) between the crystal structures of Axe2 (4JHL), 3RJT, and 5JD3.

|  | r.m.s.d (Å) |  |
| --- | --- | --- |
|  | Monomers | Dimers |
| **Axe2-3RJT** | 0.477 | 0.712 |
| **Axe2-5JD3** | 0.441 | 0.765 |
| **3RJT-5JD3** | 0.493 | 0.685 |

**Figure 1.**
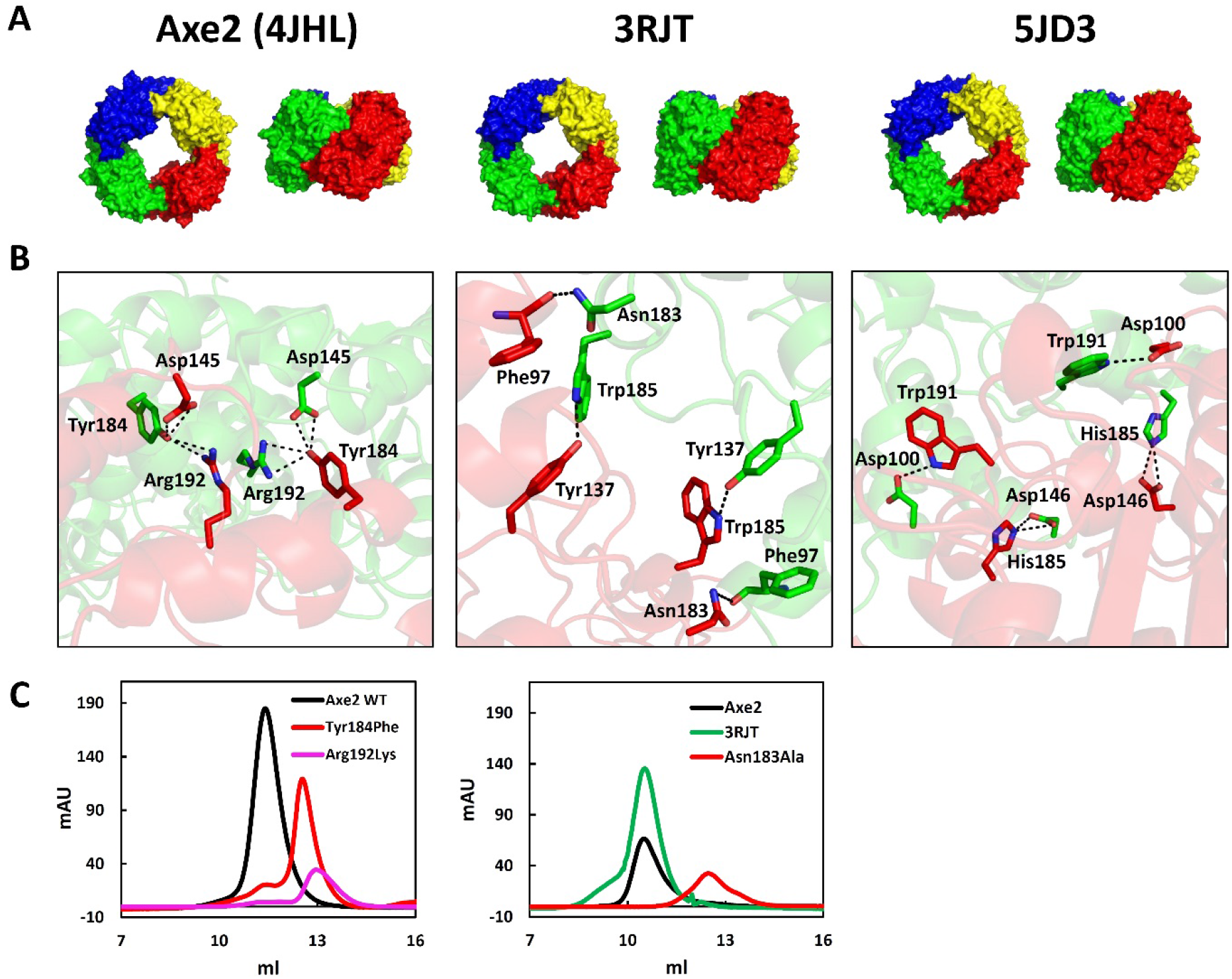
A common octameric ring structure is shared by three esterases and is maintained by sequence divergence. **(A)** The octameric structures of Axe2, 3RJT and 5JD3 are almost identical and are composed of 4 homo-dimers colored red, blue, green, and yellow. **(B)** The dimers are held together by H-bonds between different residues: Asp145, Tyr184, and Arg192 in Axe2; Phe97, Tyr137, and Asn183 in 3RJT; and Asp100, Asp146, His185 and Trp191 in 5JD3. Since the interfaces of the interacting subunits (green and red) are facing each other in isologous manner the interactions are duplicated. **(C)** Replacing Tyr184Phe and Arg192Lys in Axe2 and Asn183Ala in 3RJT resulted in proteins that were eluted as dimers based on gel filtration.

## Results

### A duplicated hydroxyl maintains the octameric structure of Axe2

The biological functional unit of Axe2 is composed of four homo-dimers, assembled in a unique doughnut-shape octameric structure (**Fig. 1A**) (19). The overall structure appears to be maintained by a combination of salt bridges, hydrogen bonds, and stacking interactions (**Fig. S1** and **Table S1**). To verify the role of key residues participating in these inter-subunit interactions, we replaced selected residues. The resulting mutants were then characterized with respect to their oligomeric state, thermostability, and catalytic activity (**Table 2**). The salt bridges between the dimers made of the conserved residues Arg55 and Glu105 are situated on interfaces with heterologous association and seems to stabilize the octameric structure of Axe2. The replacement Arg55Ala, however, did not change the oligomeric structure of the protein at 0.1 M NaCl, and only at high salt concentration, 1M NaCl, resulted in a mix population of oligomeric states. Only the more pronounced replacement Arg55Glu eluted as a dimer at 1M NaCl. Both replacements did not affect the octameric state, activity and stability at 0.1 M NaCl, indicating that the salt bridges between Arg55 and Glu105 are not crucial in maintaining the overall octameric structure of Axe2. Moreover, the characteristics of Arg55Ala and Arg55Glu mutants at different ionic strengths suggest that at low ionic strength (0.1 M NaCl) Glu105, or Glu55 (or both) are involved in altered electrostatic interactions, which may still be sufficient to maintain stability, and that higher ionic strength (1 M NaCl), contributes to elimination of all possible interactions that stabilize the octamer.

**Table 2:**
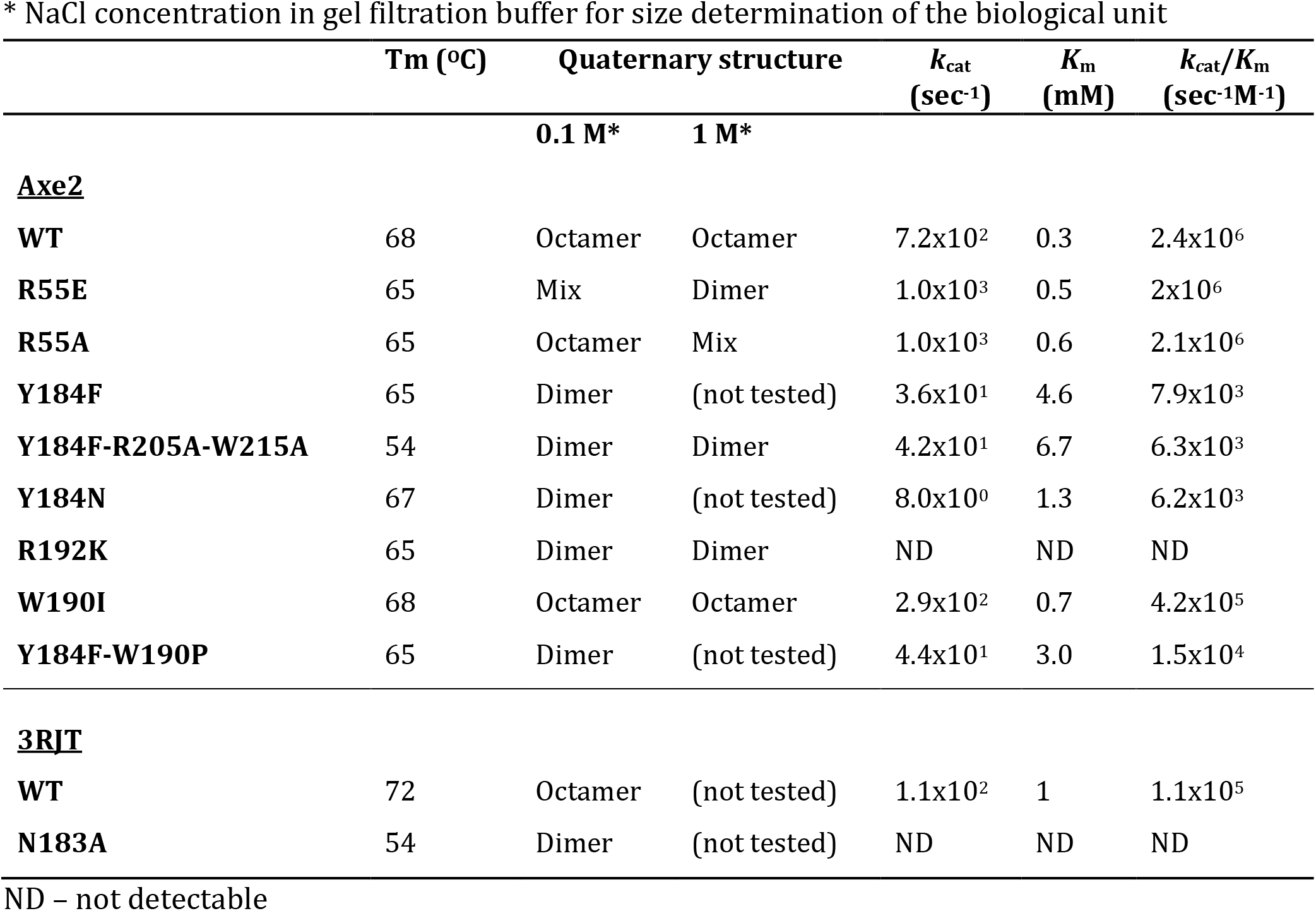
Biochemical characterization of structural mutants of Axe2 and 3RJT. The melting temperature, Tm, was determined by DSC. Reaction assay contained 200 μl of 2-naphthyl acetate (1-20 mM) in isopropanol and 600 *μ*l citrate phosphate buffer, pH 7, containing the enzyme. The experimental error for the catalytic constants was +/- 5%.

Additional interactions between the dimers are at different interfaces, which are assembled in isologous manner and comprise residues Trp95, Trp190, Tyr184, Asp145, and Arg192 (**Fig. S1**). One of the seemingly significant interactions is the π-stacking interaction between the aromatic side chains of Trp190 of one subunit and Trp95 of the other subunit. However, the Trp190Ile replacement (removing the aromatic stacking) did not affect the octameric assembly of the protein, as well as its original Tm of 68°C (crystal structure is available, PDB 4OAP). The catalytic activity of the Trp190Ile mutant did not change dramatically as well, and although the kcat was reduced from 7.2 × 10^2^ sec^−1^ to 2.9 × 10^2^ sec^−1^, this relatively small change in activity can be attributed to the fact that Trp190 is situated on the catalytic loop, thus even small local structural changes can cause relatively large fluctuations in activity. In addition, the affinity of this mutant towards the substrate remained practically the same compared to the WT. Another type of interactions at these interfaces comprises two bifurcated H-bonds that the hydroxyl of Tyr184 can form with Asp145 and Arg192 (**Fig. 1B**). These charged residues that form multiple H-bonds across the subunits are buried at the interface (**Fig. S2**) and hence represent particularly effective hotspots. Indeed, the Tyr184Phe replacement (removing the key OH group from this side chain), transformed the enzyme from octamer to a dimer (**Fig. 1C**). A similar octamer-to-dimer transformation was observed with the Arg192Lys replacement, removing the possibility for a bifurcated interaction with the Tyr184 side hydroxyl (**Fig. 1C**). Thus, the hydrogen bonds involving the side chains of Tyr184 and Arg192 are critical for the stabilization of the octameric form of Axe2. In addition, only marginal catalytic activity was observed for these two mutants in their dimeric forms, suggesting that the octameric form of Axe2 is crucial for its catalytic activity (**Table 2**).

To summarize, Axe2 represents a case where evolution exploited symmetry to stabilize the dimer interface and as a corollary, a single amino acid substitution abrogated it. The importance of symmetry in stabilizing or destabilizing an interface is further illustrated by the finding that a single mutation that abrogates the Glu105-Arg55 salt bridge at the heterologous part of the interface is less destabilizing than the elimination of the two hydroxyl groups by a single mutation at the isologous interface.

### Hydrophobic interactions maintain the dimeric structure of Axe2

As demonstrated above, a doubled hydroxyl maintains the octameric form of Axe2, and now the question was which specific interactions keep the dimer together. The two monomers are assembled in an isologous manner and appear to be held by a combination of hydrogen bonds, stacking interactions and hydrophobic interactions. Residues Arg205 and Trp215, lined up at the core of the monomers interface, create stacking interactions between the two Trp215 residues, and hydrogen bonds between the side chains of Arg205 and the backbone of Trp215 (O – NE and NE - O) (**Fig. S1**). These interactions potentially hold the C-terminal loops of the two subunits together, thereby stabilizing the dimer over the two separate monomers. Replacing these two residues (Arg205 and Trp215) to Ala, in addition to the Tyr184Phe mutant, (which was confirmed to be a dimer in all of the tested conditions) resulted in a triple mutant (Tyr184Phe-Arg205Ala-Trp215Ala). However, even in this background the protein remained in the dimer form, suggesting that the interactions between Arg205 and Trp215 are not as crucial as we suspected for the formation of the Axe2 dimer. We concluded that the dimer is held mainly by interactions between the two hydrophobic cores of the monomers, which include residues Ala39, Tyr40, Leu44, Val197 and Met201. This conclusion is reinforced by comparing the monomer-monomer and dimer-dimer interfaces. The stable monomer-monomer interface is less readily destabilized by single site mutations compared to the dimer-dimer interface despite being two-fold symmetric. Based on interface characterization (**Table 3**) the stability of the monomer-monomer interface is not attributed to larger contact area but rather to the interactions (energy) in similar sized interfaces. In fact, the dimer-dimer interactions are expected to be weaker than the monomer-monomer interactions as part of the octamer complex where they repeat 4 times. The monomer-monomer interactions, on the other hand, stand alone in the dimeric form of the protein and thus are required to be stronger.

**Table 3:** Interface characterization of Axe2, 3RJT and 5JD3. The areas of interfaces (Å^2^) are given, as well as their percentages from the total subunit area (in brackets). ΔiG is the solvation energy. Calculations were done in the Pisa server.

|  | **Monomer-monomer** |  |  | **Dimer-dimer** |  |  |
| --- | --- | --- | --- | --- | --- | --- |
| | Area<br>($\text{\AA}^2$) | $\Delta iG$<br>(kcal/mol) | Energy/area<br>(kcal/(mol $\text{\AA}^2$)) | Area<br>($\text{\AA}^2$) | $\Delta iG$<br>(kcal/mol) | energy/area<br>(kcal/(mol $\text{\AA}^2$)) |
| Axe2 | 1335.4<br>(13%) | -23.1 | 0.017 | 1258.2<br>(7%) | -11.9 | 0.009 |
| 3RJT | 947.9<br>(9.9%) | -20 | 0.021 | 1215.4<br>(7%) | -4.4 | 0.004 |
| 5JD3 | 1162.6<br>(11.7%) | -11.4 | 0.010 | 1326.5<br>(7.6%) | -15 | 0.011 |

### The octameric structures of 3RJT and 5JD3 are maintained by different interactions than Axe2

The crystal structures of two Axe2 homologues, 3RJT and 5JD3, are available and both homologues exhibit the same octameric quaternary structure as Axe2 (**Fig. 1A**). The 3RJT protein from *Alicyclobacillus acidocaldarius* subsp. Acidocaldarius DSM446, is a putative lipolytic enzyme that shares 52% sequence identity with Axe2 and contains 213 amino acids per monomer. Superposition of the octameric structures of Axe2 and 3RJT reveals a very good overlap, with an r.m.s.d. value of 0.66 Å for 212 of the 213 residues aligned. The uncharacterized 5JD3 protein was obtained from a metagenome of Lake Arreo, Spain, and shares 54% identity to Axe2 and contains 238 residues per monomer. Superposition of its crystal structure with Axe2 also shows good overlap with an r.m.s.d value of 0.61 for 206 of the 238 residues aligned. A close comparison of key residues in the crystal structures of Axe2, 3RJT and 5JD3 reveals two striking differences. First, both 3RJT and 5JD3 lack the residue equivalent to Tyr184, which was shown to be crucial for maintaining the octameric structure of Axe2. The corresponding residues at this position are Asp183 and His185, respectively. Second, the catalytic loops containing the catalytic Asp and His residues appear to be stabilized in different ways.

To characterize the 3RJT protein, we expressed in *E. coli* a synthetic gene encoding for 3RJT, and obtained the gene product. The purified 3RJT protein exhibited a Tm of 72°C and appreciable esterase activity towards 2-naphthyl acetate at 30°C, pH 7 with a *k_cat_* of 2.1 X 10^2^ sec ^-1^ (**Table 2**). The calculated molecular weight of 3RJT is 26,415 Da, and based on gel filtration, the protein is about 200,000 Da, corresponding to an octameric structure. The salt bridges between the dimers in Axe2 also exist in 3RJT, with the corresponding residues Arg54 and Glu104. The main interactions between the dimeric subunits in 3RJT are positioned as in Axe2. However, 3RJT appears to utilize different residues than Axe2 for stabilizing the octamer since it lacks the corresponding crucial Tyr184 residue of Axe2. In 3RJT, the side chains of Trp185 (NE1) and Asn183 (ND2) interact with Tyr137 (OH) and the backbone of Phe97 (O), respectively (**Fig. 1B**, middle). Indeed, the replacement Asn183Ala produced a dimer of the 3RJT protein (**Fig. 1C**), which, unlike the Axe2 dimer, had a pronounced effect on the Tm of the protein, reducing it from 72°C for the WT, to 54°C for the dimer. This dimeric form render the protein inactive (**Table 2**), emphasizing the importance of the octameric configuration for enzyme catalysis, as in the case of Axe2. As with Axe2, the monomers of the 3RJT dimer are held by hydrophobic interactions between residues Leu39, Ala42, leu196, and Leu200. Similarly to Axe2, the energy gain on complex formation is higher for the monomer-monomer interface than for the dimer-dimer interface. However, unlike Axe2, the 3RJT monomer-monomer interface is smaller by 22% than the dimer-dimer interface, nevertheless, the energy to area ratio for Axe2 monomer-monomer interface is 0.017 and for 3RJT is 0.021 kcal/(mol Å^2^), which makes the 3RJT dimer even slightly more stable than Axe2 (**Table 3**).

The quaternary structure of the 5JD3 protein (obtained from a metagenome of Lake Arreo, Spain) is similar to the octameric ring of Axe2 and 3RJT. The interfaces associated in a heterologous manner comprise the conserved salt bridges, involving residues Arg56 and Asp106 in 5JD3, as in Axe2 and 3RJT. The interfaces that are critical for the association of the dimers comprise His185, which corresponds to the critical Tyr184 and Asn183 residues in Axe2 and 3RJT, respectively. The 5JD3 octamer is most likely to be maintained by the interactions between the side chains of His185 (NE2) with Asp146 (OD2), and Trp191 (NE1) with Asp100 (OD2) (**Fig. 1B**, right). Thus, 5JD3 demonstrate a third arrangement of maintaining the octameric structure in the Axe2 family. Similarly to Axe2 and 3RJT, the 5JD3 dimer is held by the hydrophobic core of its monomers with residues Ala44, Leu41 and Leu198. Unlike Axe2 and 3RJT the area to energy ratio in 5JD3 is the same for the monomer-monomer and dimer-dimer interfaces and is 0.01 kcal/(mol Å^2^) (**Table 3**). Thus, in the case of 5JD3 there is no prevailing effect of either the area size or the type of interactions on the stability of the interfaces, and it seems that the dimer and the octamer are stable to the same extent.

### Different interactions stabilize the catalytic loop in the three Axe2 homologous

Based on the crystal structures of the three Axe2 homologous the catalytic loops of Axe2, 3RJT and 5JD3 appear to be stabilized by different interactions (**Fig. 2**). The catalytic loop of Axe2 is stabilized by pro195, H-bond between Arg192 and Tyr184 from the adjacent subunit, and stacking interactions involving Trp190 and Trp95 from the adjacent subunit. The 3RJT protein does not have the corresponding residues and its catalytic loop is stabilized by two Pro residues (Pro189 and Pro194), located on each side of the loop, and by a hydrogen bond between Arg191 and Glu139 in the middle of the loop. It seems that in the case of 3RJT, there are no inter-subunit interactions that stabilize the loop as in the case of Axe2. Similarly to Axe2 and 3RJT, 5JD3 uses Pro196 on one side of the loop and the other side of the loop is stabilized by a hydrogen bond between Trp191 (as in Axe2) and Asp100. The middle of the catalytic loop is stabilized in 5JD3 by interaction between Arg193 and Glu141 as with shares 3RJT.

**Figure 2.**
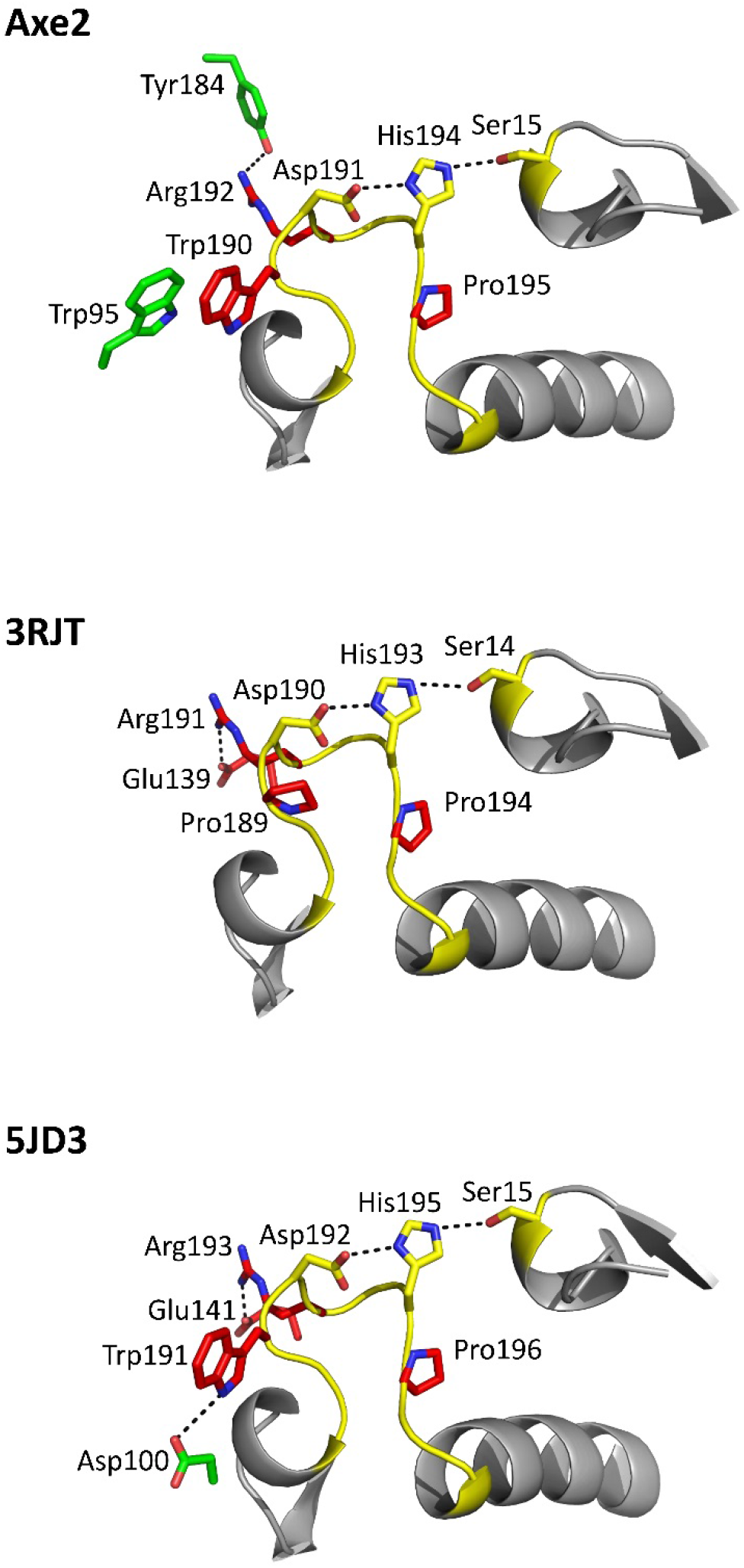
The conserved catalytic loop is stabilized by different interactions in Axe2, 3RJT and 5JD3. The catalytic loop (yellow) comprises the catalytic His and Asp is in close proximity to the nucleophilic Ser (yellow). The three homologues use different amino acids to stabilize the loop: Axe2 uses intermolecular stacking interactions (Trp190 and Trp95) and H-bonds (Arg192 and Tyr184), as well as Pro195 residue; 3RJT uses 2 Pro residues in either side of the loop (Pro189 and Pro194) and intramolecular H-bond in the middle (Arg191 and Glu139); and 5JD3 uses intermolecular H-bond (Trp191 and Asp100), intramolecular H-bond (Arg193 and Glu141), and Pro196.

### The interfaces of the Axe2 homologues are not conserved relatively to their surfaces

The three homologous, Axe2, 3RJT, and 5JD3, provide an example of divergence of structurally important interactions that stabilize the octameric assemblies of the proteins. This is intriguing since critical residues are usually conserved and are part of conserved interfaces. We thus addressed the question whether the monomer-monomer and dimer-dimer interfaces are conserved in comparison to the general surface of the proteins. To do this analysis, we constructed surface patches having the same area as the interfaces in the three homologous and then obtained conservation scores for every patch and compared the surface scores to the interface score (**Fig. 3**). Surprisingly, in all cases, the monomer-monomer and dimer-dimer interactions of the three homologues did not appear to be conserved relative to other protein surfaces of the same surface area.

**Figure 3.**
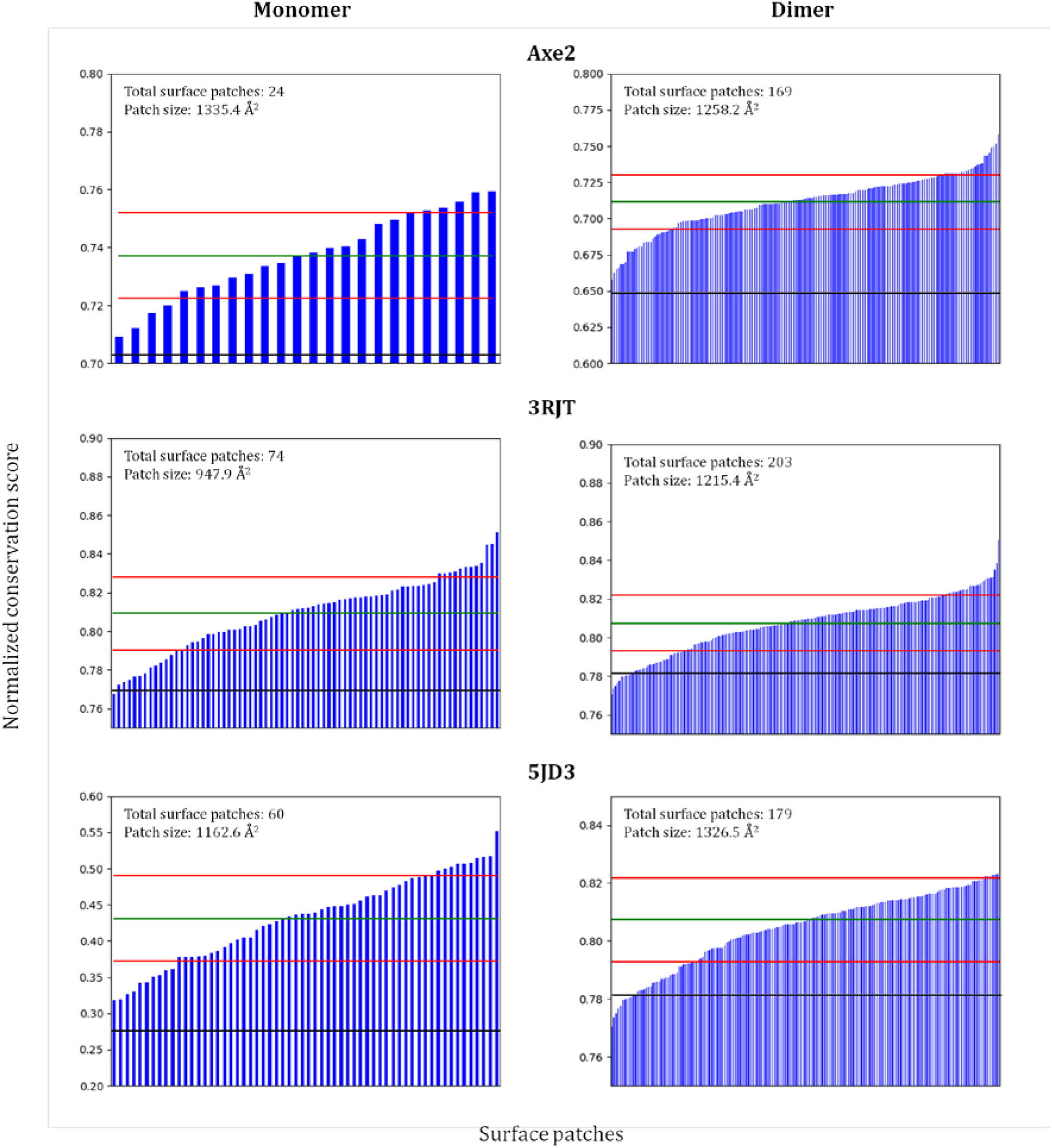
Axe2, 3RJT and 5JD3 interfaces are not conserved relatively to the surfaces. Surface patches equal in size to the interface patch were constructed, and their conservation scores compared to the interface patch. The surface patches do not contain amino acids that belong to the interfaces. The interface conservation score is represented by a black line, and is well below the conservation average of all the surface patches, represented by the green line. Red lines – standard deviations of surface conservations. Interface area calculations were done in the Pisa server and conservation scores were calculated using the Consurf server.

### Additional Axe2 homologues share the octameric structre

The fact that Axe2, 3RJT and 5JD3 share similar ring-shaped octamer structure, yet different types of interactions maintain their quaternary assemblies, prompted us to investigate the quaternary structures of additional homologous proteins in the same family. For such study, the idea was to select homologues that have relative lower degree of sequence identity, specifically in the potential interface regions between the subunits. With this objective in mind, we constructed a phylogenetic tree and selected five potential proteins which are homologous to Axe2, but which are located on different branches of the tree and contain different type of residues at the critical interface positions (**Fig. 4A**). The proteins selected are termed Pjd, Pal, Pda, Sku and Cph. Based on the primary sequences of the selected proteins, we obtained the corresponding synthetic genes, and produced the proteins in *E. coli* (see supplementary material). Since the proteins were not characterized previously and had no available structures, we used a number of experimental techniques to verify their functional molecular weight, oligomeric state, general structure, and esterase activity towards the standard substrate 2-naph-OAc. Based on gel filtration, all of the homologues appear to have molecular weights of about 200,000 Da, indicating octamer assemblies; in addition, DLS studies confirmed that they all share similar hydrodynamic diameters in solution corresponding to circular octamers (**Table 4**). TEM analysis revealed that all proteins had ring structures similar to Axe2 and 3RJT (**Fig. 4B**), confirming the octameric assemblies. In addition, all of the proteins exhibited esterase activity.

**Table 4:** Size characterization of Axe2 homologues. Axe2 is given at the top for comparison. The molecular weights (Mw) were determined by gel filtration and the number of subunits was estimated from the Mw. The diameters of the proteins were determined by dynamic light scattering (DLS). All the proteins were active on 2-Naph-OAc.

| Short name | UniProt/Uniparc number | Protein length | Mw | Estimated Number of subunits | Hydrodynamic Diameter at DLS (nm) |
| --- | --- | --- | --- | --- | --- |
| Axe2 | Q09LX1 | 219 | 194,341 | 8 | 12.3 |
| 3RJT | C8WT89 | 216 | 221,960 | 8 | 11.8 |
| Pjd | C6D722 | 220 | 178,663 | 8 | 11.7 |
| Pal | K4ZJU9 | 213 | 209,053 | 8 | 11.3 |
| Pda | UPI0003713BD1 | 218 | 192,371 | 8 | 10.4 |
| Sku | UPI00037F5CD6 | 222 | 211,971 | 8 | 12.4 |
| Cph | A9KJ38 | 219 | 189,723 | 8 | 12.5 |

**Figure 4.**
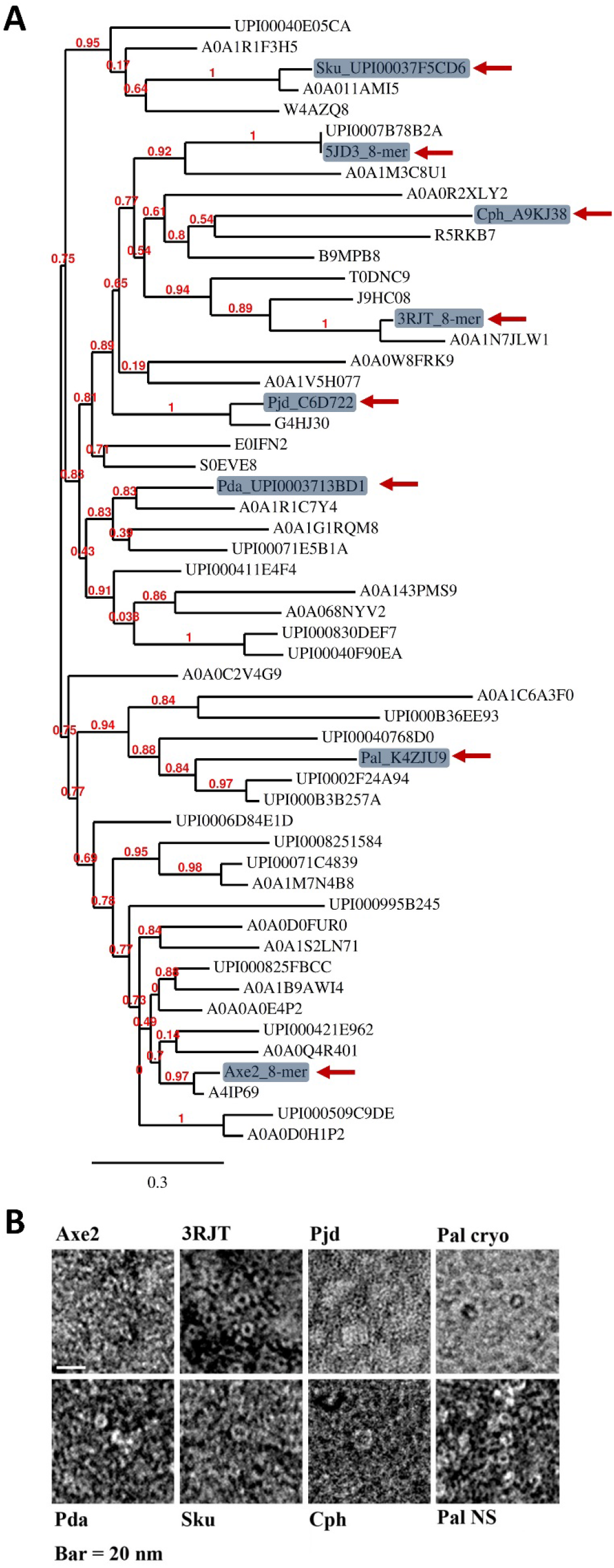
Other members of the Axe2 family share the common octameric ring structure. (**A**) Phylogenetic tree of Axe2 and its homologues. Sequences obtained through the ConSurf server ^46^ (MSA was built using MAFFT, The Homologues were collected from UNIREF90, homolog search algorithm: HMMER, HMMER E-value: 0.0001, 1 iteration. 54 sequences were selected with E-value of between 6e-145 and 1.9e-62). The phylogenetic tree was generated using the phylogeny.fr server (55). (**B**) Homologues that were chosen from different branches of the phylogenetic tree have similar ring structures as demonstrated in TEM images. These structures were confirmed by gel filtration to be octamers (see table 2).

### The common octamers are maintained by non-conserved interactions

Based on crystal structures and biochemical studies of Axe2, 3RJT, and 5JD3, we showed that a specific residue (corresponding to Tyr184 in Axe2) is critical for maintaining the octameric structures of the proteins. Since 3RJT has Asn and 5JD3 has His at this position, we concluded that the octameric ring is maintained by non-conserved interactions. This conclusion is reinforced by the characterization of additional five homologues, which were shown to be octameric rings, and by the fact that the critical position is not conserved in the Axe2 family (**Fig. S2**). Overall we observed four different residues in this position among the homologues we investigated: Tyr (in Axe2, Pda, and Pal), Asp (in 3RJT), His (in 5JD3, Pjd, and Cph) and Ile (in Sku).

As mentioned above, the dimers of Axe2, 3RJT, and 5JD3 are maintained by hydrophobic interactions between the monomers. Each protein has a different set of residues for this purpose. We used modeling techniques in order to identify the potential interfaces and specific residues that maintain the dimers in the other homologues. Our main conclusions were obtained from rigid body docking (PyDock), which were followed by Normalized Interface Propensity (NIP) (see supplementary material for method details, and **Fig. S3**). The residues with significant NIP values define potential binding sites. We first built the monomeric models for each of the homologues. Then, based on these models we determined the potential binding sites between the monomers, which allowed us to construct the most probable dimers. All the homologues have the same monomer-monomer interacting interfaces to construct the dimeric subunit and they all contain a cluster of non-conserved hydrophobic residues, potentially holding the two monomers together (**Table 5**). These non-conserved interactions in non-conserved interfaces (**Fig. S4**) demonstrate clearly sequence divergence that maintains a favorable dimeric arrangement. Taken together, our analysis indicates that the common octameric structure in this new group of esterases is maintained by non-conserved interactions, demonstrating sequence divergence at the level of quaternary structure.

**Table 5:**
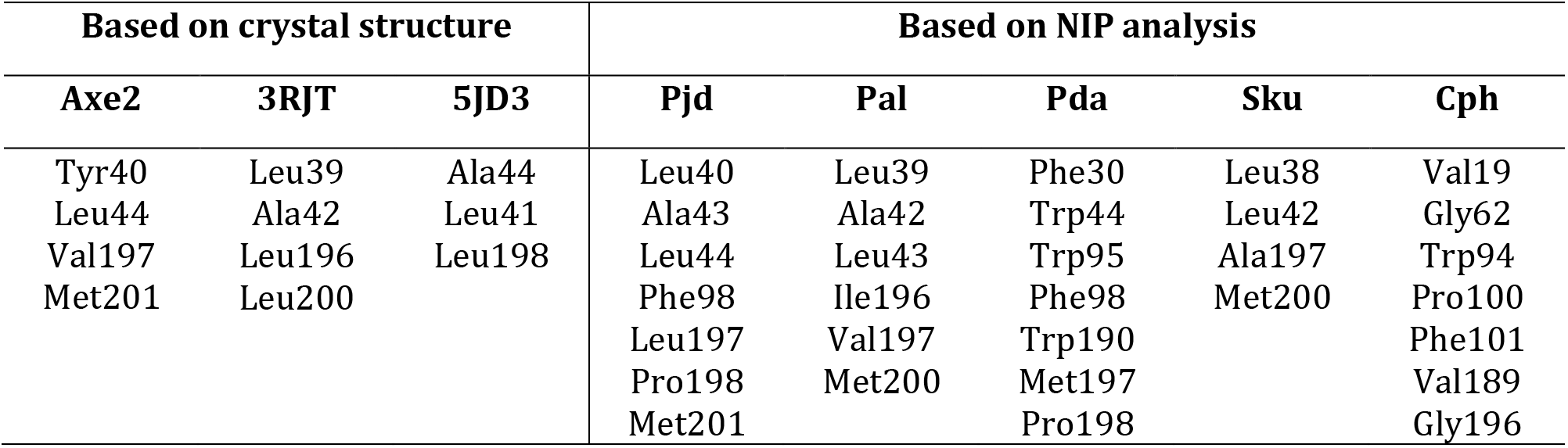
Residues of the hydrophobic core of Axe2 homologues. The cores maintain the dimeric subunits of the proteins. Residues were detrmined by either the available crystal structures or by NIP analysis (>0.2).

## Discussion

### The octameric structure is essential for the activity of Axe2 and 3RJT

In both proteins, Axe2 and 3RJT, the dimeric forms were practically inactive, revealing the importance of the octameric assembly for the catalytic activity. Oligomeric state influencing enzyme activity was reported before, for example, where conformational changes in the monomer resulted in reduced activity compared with the dimer (23). Here, we suggest that oligomerization affects the activity of the enzymes by either stabilizing the active sites or/and creating a favorable charge around them. The catalytic triad Ser-His-Asp is located at the interface between the two dimers. Based on the crystal structures and our results, the octameric form stabilizes the catalytic loops of Axe2, 3RJT, and 5JD3 in different ways (**Fig. 2**). An attempt to stabilize the catalytic loop of the dimeric Axe2, Tyr184Phe, by further replacing Trp190 with Pro (i.e. Tyr184Phe-Trp190Pro, whose crystal structure was recently determined (24)) did not resulted in improved activity, further supporting the notion that oligomerization is crucial for maintaining the catalytic activity. It is also possible that the octameric structure provides an overall negative charge around the active sites in Axe2 (**Fig. S5**). The negative charge may facilitate expulsion of the product (the acetate) from the active site in the second catalytic step, and thus, reduces product inhibition (25), and maintains the optimal pH of the enzyme at 8.5 (15), similarly to the pH dependence protection in arylsulfatase (26) (27). Indeed, we predicted that the rate-limiting step in Axe2 is the second deacetylating step in which acetate is removed from the enzyme (15). Negative potential at the active site was also observed in glycoside hydrolases, and has been connected to substrate binding (28) (29) (30).

### The octameric assembly is a common ancestor of the Axe2 family

In this study we showed that members of the Axe2 family share common quaternary octameric structures which are maintained by non-conserved interactions, resulting from sequence divergence. Replacements of either Tyr184 or Arg192 in Axe2, and Asn183 in 3RJT, resulted in the dimeric forms of the proteins. Previous studies have shown that during evolution, the switch between “surface patch” to “interface patch” is usually satisfied by changing only two amino acids per 1000 Å^2^ (31). In another study of artificial design of new oligomers by amino acid substitution it was shown that some proteins needed only a single amino acid replacement to associate to a higher-order complex (32). In our case, a similar phenomenon has been observed, as a single amino acid replacement in an isologous associating interfaces switched the interface patch into a surface patch, in both Axe2 and 3RJT. In the case of monomer-monomer interactions, our NIP analysis showed mainly non-conserved hydrophobic residues, potentially involved in formation of the dimers. Interestingly, hydrophobic interactions were shown to be more common in protein folding rather than protein-protein interactions (33). Similar structures stabilized by different interactions (resulting from sequence divergence) have been observed before at the tertiary structural level of protein folding. For example, the third domain of ovomucoid and the C-terminal fragment of ribosomal L7/L12 protein (CTF) share a similar fold, which is stabilized by different interactions in the two proteins (34). The globin family is an example of a very large diverse family with a common fold (35) (36) (37), whereas the ubiquitously TIM-barrel fold is shared among proteins with very low sequence identity belonging to different protein families (38) (39). The fact that very different sequences give rise to the same fold suggests that this fold is of high intrinsic stability (40). In contrast, sequence divergence in protein families usually leads to different quaternary structures (which may reflect evolutionary steps) among the group members (41) (42). Such divergence can also lead to a shift of subunits to other regions of the protein surface (43). Similarly, proteins with conserved quaternary structures usually have conserved interfacial residues compared with the rest of the surface (44). In this respect, Axe2 and its homologues reveal a unique phenomenon where a conserved quaternary structure is stabilized by non-conserved interactions among homologous proteins. We therefore suggest that proteins in the Axe2 family of esterases, represented originally by Axe2, have evolved from a common stable octameric ancestor. This favorable quaternary structure was maintained throughout evolution by a set of divergent interactions that increased their stability at different environmental conditions, or changed their substrate specificity. Further analysis is required to determine whether the quaternary structures of other proteins with lower sequence identity and similar fold to Axe2 family are a consequence of further sequence divergence (i.e. originating from the octamer) or are evolutionary steps in the formation of the octameric structure, as stochastic transitions (due to sequential mutations and genetic drift) lead to various multimeric configurations at both directions (7).

## Materials and Methods

### Enzyme overexpression, mutagenesis and purification

Protein expression and purification of Axe2 (GenBank accession number ABI49953.1) were performed essentially as previously described ^15^. Site-directed mutagenesis was performed using the QuikChange site-directed mutagenesis kit (Stratagene). The heat treatment step for the Axe2 mutants Trp215Ala, Arg215Ala-Trp215Ala, and Tyr184Phe-Arg215Ala-Trp215Ala was milder, 40°C for 20 min. With the Axe2 Tyr184Phe and Trp190Pro mutants, the heat treatment step was omitted and a protamine sulfate step (0.1% final concentration) was added to remove nucleic acids prior to the gel filtration step. The lipolytic Axe2 homologue from *Alicyclobacillus acidocaldarius* (NCBI reference sequence: YP_003185992.1; referred here as 3RJT), as well as its Asn183Ala mutant, were produced and purified as for Axe2 WT.

### Cloning, expression and purification of Axe2 homologous genes

The synthetic homologous genes were ordered from Integrated DNA Technologies-IDT (Iowa, USA) cloned into pET9d (for more details see supplementary). Overexpression and purification are as mentioned above, only with a protamine sulfate step (0.2% final concentration) and no heat treatment. To improve protein solubility the cultures were grown overnight at 18°C after one hour of incubation at 37°C. Typical final OD_600_ values for the overnight TB cultures were 7–24.

### Biochemical characterization

The molecular weights of native proteins in solution were determined by gel filtration using Superpose 12 or Superdex 200 10/300 filtration columns (GE Healthcare Life sciences) running at 0.5 ml/min with 50 mM Tris-Cl buffer, pH 7, containing either 0.1 or 1 M NaCl and 0.02% sodium azide. Esterase activity was measured using 2-naphthyl acetate as a substrate as previously described ^15^. Melting temperature (Tm) was determined using a MicroCal VP-DSC differential scanning calorimeter (MicroCal Inc., Northampton, MA), with protein samples of 0.3-0.5 mg/ml. The hydrodynamic diameter of the proteins was evaluated by dynamic light scattering (DLS, Cordouan Technologies, VASCO^™^ - 2 Particles Size Analyzer), equipped with a 65 mW diode laser, operating at 658 nm wavelength at a scattering angle of 135°. Samples were measured at 25°C at the following concentrations: Axe2, 2mg/ml; Pal, 5.8 mg/ml; Cph, 4.3 mg/ml; Pda, 3.5 mg/ml; Sku, 3.6 mg/ml; and Pjd, 4.4 mg/ml. The results were analyzed with the nanoQ software.

### Interface characterization and conservation analysis

All calculations, parsing and structure manipulations were done using Biopython (45), pandas (46) and Numpy (47). PDB structure files of Axe2, 3RJT and 5JD3 were retrieved from the PDB web server (48). Accessible solvent area (ASA) and residues involved in the interfaces between complex subunits were calculated using PISA web server (49) for the monomers and for the dimers assemblies, while conservation scores were calculated using the Consurf web server (50). To determine interface and surface patches conservation we extracted the calculated ASA (Å^2^) of the interface between subunits, and used this area as a baseline to calculate all the patches with the same radius (thus, area (Å^2^)) on the protein surface. These patches did not consist any residue in common with the interface patch. For each patch, a mean conservation score was calculated based on the Consurf conservation results, following normalization.

### Protein modeling and docking analysis

Homologues models were constructed and analyzed using the Phyre2 web-server (51). Conservation calculations were done using the ConSurf server (52). All docking calculations were done with the PyDock web-server (53). Normalized Interface Propensity (NIP) calculations were performed using the local version of the PyDock 3.0 package (54). The docking results were re-scored with the novel democratic rescoring method using the pyDockRescoring web-server.

### Data Availability

All data generated or analyzed during this study are included in this published article (and its Supplementary Information files). The datasets generated during and/or analysed during the current study are available from the corresponding author on reasonable request.

## Acknowledgments

This work was supported by the Israel Science Foundation Grants 500/10, 152/11 and 1505/15, the I-CORE Program of the Planning and Budgeting Committee, the Ministry of Environmental Protection, and the Grand Technion Energy Program (GTEP), and comprises part of The Leona M. and Harry B. Helmsley Charitable Trust reports on Alternative Energy series of the Technion, Israel Institute of Technology, and the Weizmann Institute of Science. Y.S. acknowledges partial support by the Russell Berrie Nanotechnology Institute and The Lorry I. Lokey Interdisciplinary Center for Life Science and Engineering, Technion. Y.S. holds the Erwin and Rosl Pollak Chair in Biotechnology at the Technion. We thank Dr. Inbal Abutbul-Ionita and Dr. Ludmila Abezgauz for the TEM analysis, Noam Adir and Yael Pazy Benhar for critical reading of the manuscript and for the useful suggestions of the anonymous reviewers.

## Author Contributions Statement

O.A., G.S., and Y.S. designed the study; O.A., R.S., N.L., and S.L. performed the experiments; O.T. performed the modelling and docking. All authors analyzed the results and contributed to writing and editing of the manuscript.

